# The neural basis of unfamiliar face recognition in humans with fMRI frequency-tagging

**DOI:** 10.64898/2026.11.12.764897

**Authors:** Marie-Alphée Laurent, Jean-Baptiste Durand, Xiaoqing Gao, Nathalie Vayssière, Isabelle Berry, Bruno Rossion, Benoit R. Cottereau

## Abstract

Faces provide the most diagnostic information for identifying individuals in humans. Understanding the neural mechanisms involved in face identity recognition (FIR) is particularly important for unfamiliar faces, for which identity cannot rely on encoded multimodal semantic associations. Here, we investigated the neural basis of FIR using an original frequency-tagging fMRI approach providing objective, sensitive, and reliable measures of face recognition. Ten human participants were presented within a rapid 6 Hz stream of natural images of a single unfamiliar identity in two conditions: (1) with the same face image across low-level changes (size, luminance, contrast) only, or (2) with images varying in background, size, head orientation, gaze direction, etc. Every 9 s during a 243 s run, seven images of different unfamiliar identities were introduced in bursts. For each participant and each condition, we recorded runs with upright and inverted faces. Analyses were performed within face-selective regions (defined from a frequency-tagging localizer) and in the Fourier domain, where individual face discrimination responses were objectively identified and quantified at the identity-change frequency (0.111 Hz). We observed robust image-based individual face discrimination responses across both conditions in core face-selective ventral regions (LatMidFG and IOG). In contrast, responses in early visual cortex and the pSTS were negligible in our second condition, which required generalization across viewpoint variations. In addition, the LatMidFG showed a reliable inversion effect, confirming that the observed responses cannot be attributed solely to low-level image differences. Overall, these findings highlight the value of fMRI frequency-tagging for objectively characterizing the cortical network underlying unfamiliar FIR, providing evidence for high-level discrimination and generalization of unfamiliar face identity in the human ventral occipito-temporal face-selective network.

## 1. Introduction

Faces play a central role in human social interactions, providing the most diagnostic signals for identifying individuals in our species (Sheehan and Nachman, 2014). Face Identity Recognition (FIR) entails a behavioral or neural response that is both distinct from responses to other facial identities (i.e., discrimination) and sufficiently reliable across different views of the same individual (i.e., generalization). Given the constant exposure to novel faces in modern human societies, the ability to discriminate and generalize identity for faces that have not been encoded in memory (i.e., unfamiliar) is a key human function. Hence, although long-term familiarity with facial identities undoubtedly improves FIR (Bruce, 1982; Bruce et al, 2001; Johnston and Edmonds, 2009), neurotypical human adults are able to recognize (i.e., discriminate and generalize across) exemplars of unfamiliar faces rapidly (Jacques et al., 2007), automatically (Liu-Shang et al., 2014; Yan et al., 2019), and with high accuracy (Rossion and Michel, 2018). FIR also works under head rotation and lighting changes (Favelle et al., 2017; Rossion and Michel, 2018) and degraded viewing conditions (Bruce et al., 2001; Gilad-Gutnick et al., 2012; Bindemann et al., 2013). Moreover, since unfamiliar faces are devoid of associated semantic and affective cues facilitating recognition, understanding the neural basis of unfamiliar FIR is fundamental for understanding FIR based on visual cues and memory (Rossion, 2018).

Lesion studies in patients with prosopagnosia (e.g., Sergent and Signoret, 1992; Barton, 2008), transcranial magnetic stimulation (TMS; Pitcher et al., 2007), and transient impairments during focal intracerebral stimulation (Jonas et al., 2012; Volfart et al., 2023) have consistently demonstrated the critical involvement of the ventral occipito-temporal cortex (VOTC) in FIR. Intracranial electroencephalographic (iEEG) recordings at a macroscopic level of organization have further supported the central role of the VOTC in FIR and highlighted the right-hemispheric dominance of this function (Jacques et al., 2020).

However, the more numerous neuroimaging studies, mainly with functional magnetic resonance imaging (fMRI), that have attempted to provide neural maps of FIR with pictures of unfamiliar faces have reported more inconsistent results. On the one hand, face-selective regions in the VOTC activated by unfamiliar faces, such as the lateral portions of the inferior occipital gyrus (IOG or ‘Occipital Face Area’, OFA) and the middle fusiform gyrus (MidFG or ‘Fusiform Face Area’, FFA) (Sergent et al., 1992; Haxby et al., 2000; Rossion et al., 2012; Zhen et al., 2015; Grill-Spector et al., 2017) have been reported to be sensitive to immediate unfamiliar face identity changes (fMRI-adaptation; e.g., Gauthier et al., 2000; Grill-Spector and Malach, 2001; Davies-Thompson et al., 2009; Gentile and Rossion, 2014). On the other hand, fMRI studies investigating response patterns evoked by different individual faces across multiple voxels (Multivariate Pattern Analysis or MVPA) have shown rather negative or more mixed results in these posterior face-selective regions, instead emphasizing the contribution of anterior temporal lobe regions (Kriegeskorte et al., 2007; Natu et al., 2010; Nestor et al., 2011; Goesaert and Op de Beeck, 2013; Anzelotti et al., 2013; Anzelotti and Caramazza, 2014; Anzellotti and Caramazza, 2017). In addition, proponents of the MVPA approach have criticized fMRI-adaptation approaches of lacking specificity (i.e., detecting all kinds of changes beyond face identity; Mur et al., 2010), while at the same time showing visual discrimination based on a restricted set of identities, typically presented in non-ecological contexts (e.g., cropped faces, artificial backgrounds), providing limited evidence for generalization across image variations (e.g., Nestor et al., 2011; Anzellotti and Caramazza, 2014).

Here we address these issues by measuring unfamiliar neural FIR activity in humans with an fMRI-adaptation paradigm. To this end, we capitalized on an original frequency-tagging approach adapted for fMRI (Gao et al., 2018; Laurent et al., 2023), which offers highly sensitive, reliable, and objective measures of face-selective responses, while benefiting from the high spatial resolution and extensive spatial coverage of fMRI. Specifically, we extended a validated face individuation frequency-tagging paradigm in (i)EEG (e.g., Liu-Shang et al., 2014; Jacques et al., 2020; Rossion et al., 2020 for review) to fMRI, using a large pool of natural face images to increase ecological validity (Figure 1). In this paradigm, the same unfamiliar face identity is presented repeatedly at a high frequency rate of 6 Hz, allowing viewers only one fixation per face/image, either across minimal low-level (i.e., size, global luminance and contrast) changes (condition 1) or substantial changes in view, involving both low-and high-level changes in background, head orientation, and gaze direction (condition 2). Every 9 seconds (0.111 Hz), a ‘mini-burst’ of 7 new unfamiliar face identities alternating with the repeated identity is introduced (Figure 1). This paradigm takes advantage of both the optimal frequency rate for face individuation at 6 Hz (Alonso-Prieto et al., 2013; Gentile & Rossion, 2014; Retter et al., 2021) and of neural adaptation to a repeated face identity. The periodic (0.111 Hz) introduction of new unfamiliar face identities should elicit a rebound in the blood oxygenation level dependent (BOLD) response in a neural population that discriminates between facial identities, leading to an objectively identifiable 0.111 Hz BOLD signal in the frequency spectrum. With this approach, including further low-level control with picture-plane inversion, we report robust evidence of FIR responses to unfamiliar faces across the human core face-selective network, particularly in its ventral component for high generalization.

**Figure 1.**
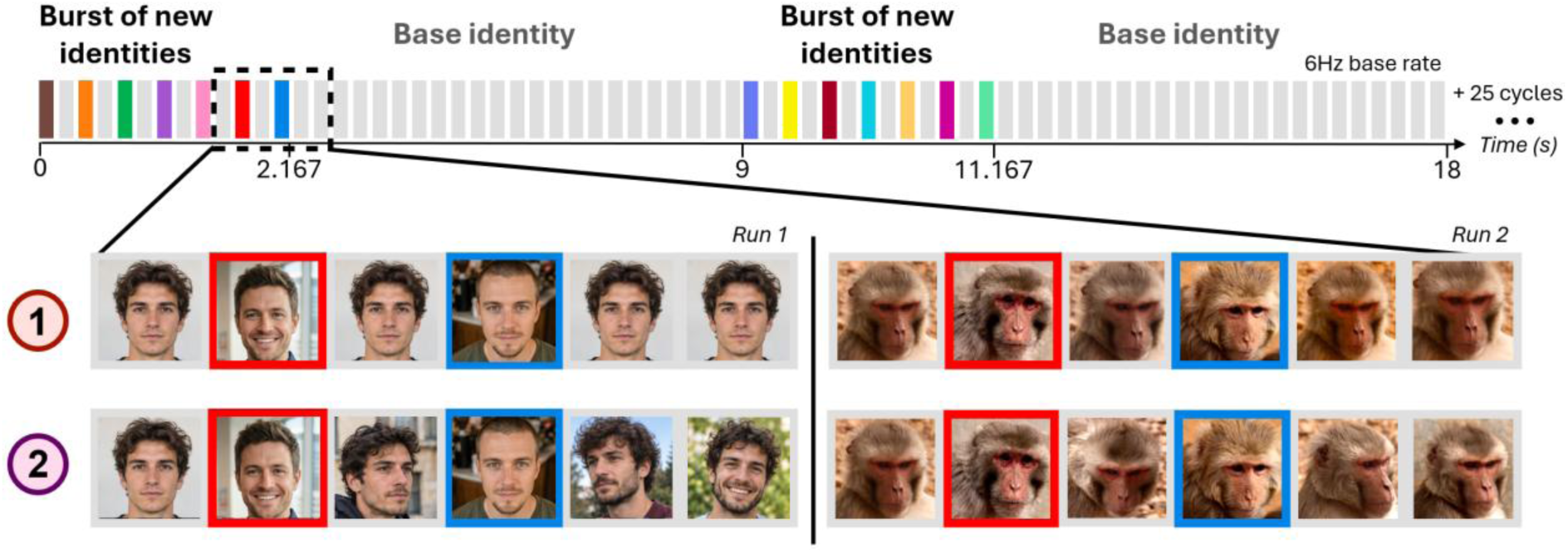
Illustration of the Face Individuation paradigm. The protocol relies on a rapid periodic presentation of visual stimuli at 6 Hz. Each 9-second cycle starts with a 2.167 seconds “burst” of stimuli from the alternate category of individuals. This burst contains seven different unfamiliar identities (illustrated with colored frames) alternating with six images of the base facial identity (illustrated with grey frames). The single base identity is presented for the rest of the cycle in two conditions: (1) the same face image, or (2) different images and viewpoints. Each run contains 27 cycles and lasts 243 seconds. Note that within each run, we presented faces from only one primate species (human or macaque), and faces were shown upright or inverted in separate runs. Human faces shown in this figure are AI-generated synthetic images and do not depict real individuals. Real photographs of human faces were used in the experiments.

## 2. Methods

### 2.1. Participants

Ten healthy individuals (6 females; mean age: 24.5 ± 2.2 years; 9 right-handed) were included in this study. All participants had normal or corrected-to-normal vision and none reported having a history of neurological or psychiatric disorder. Prior to fMRI scanning, all participants performed the computerized version of the Benton Face Recognition Test (BFRT-c, Rossion and Michel, 2018, see Benton and Van Allen, 1968 for the original version of the test) to assess their face discrimination abilities. All participants performed within a normal range, with a mean score of 46/54 (± 4.1) (range 39-52) and a mean response time (RT) of 160.9 ± 52.9 sec (range 80-232). They all gave written informed consent to participate to the study and the protocol was approved by the French Ethics Committee of the CPP Sud-Ouest Outre-Mer IV (N°2015-A01801-48).

### 2.2. Visual stimuli

Two sets of visual stimuli were used in this study. In the face categorization paradigm, stimuli included natural color images of 100 distinct, non-famous human faces, 89 rhesus macaque face images, and 195 non-face object images (from Gao et al., 2018 and Laurent et al., 2023). Each face image contained only a single human or monkey face.

Stimuli used in the face individuation paradigm were selected from a pool of natural, unsegmented color images containing 44 human and 44 monkey faces (Supplementary Figure 1). Among these sets of images, 12 different male identities from each category were presented across 6 different viewpoints as baseline identities in distinct sequences (Supplementary Figure 2), while the other 25 identities were presented as alternate identities (Figure 1). The face images were selected from publicly available image sources, and the 12 monkey face images with variations originate from a dataset created by Taubert and colleagues at the National Institute of Mental Health in Bethesda, USA. After each scanning session, none of the participants reported recognizing any of the human faces. Face images varied significantly in size, lighting conditions, color, 3D orientation, and background, thereby minimizing the contribution of low-level cues to face discrimination neural responses and ensuring that these responses reflect generalization across variable exemplars (Rossion et al., 2018). To further control for the potential contribution of low-level visual cues to category-selective responses, the stimuli were resized to 256 x 256 pixels and equalized for luminance and contrast (average values of 115 ± 0.4 and 0.55 ± 0.06 were used). Inverted versions of these faces were generated by vertically flipping each image.

### 2.3. Scanning procedures

To localize brain regions responding more to faces than to non-face objects, each participant was first tested using a validated frequency-tagging face categorization paradigm (Gao et al., 2018 and Laurent et al., 2023). During a run, images of non-face objects were displayed in random order on the screen at a rate of 6 Hz (i.e., 6 images/sec) for 243 seconds (162 TRs). Every 9 seconds (i.e., at F = 1/9 = 0.111 Hz), seven randomly selected images from the face category (human faces or monkey faces) were presented, alternating with six images of non-face objects for 2.167 seconds.

In the face individuation paradigm (Figure 1), natural images of a single unfamiliar identity (in grey) were presented within a rapid 6 Hz stream (i.e., 6 images/sec, ensuring one gaze fixation per image) in two conditions. In the first condition, the same face image was presented across low-level changes (± 10% variations in size, luminance, contrast) only. In the second condition, different images of the same identity were shown, introducing additional changes (e.g., background, size, position, head orientation, gaze direction, etc.). Every 9 seconds (i.e., F = 1/9 = 0.111 Hz) during a 243-s run, seven randomly selected images of different unfamiliar identities were introduced in bursts (in colored frames). Importantly, these new face identities alternated with six images of the base facial identity for 2.167 seconds to avoid neural adaptation and maximize the contrast between the two identities. Thus, as illustrated in Figure 1, despite controlling for low-level variations (size, luminance, contrast), changes of identity in condition 1 are *unspecific*, being confounded by changes in head position, orientation, background, relative size, etc. In contrast, in condition 2, variability in head position, orientation, background, relative size, etc. are introduced also in the repeated face identity, controlling for these factors by variation (rather than normalization). In both conditions, faces were presented upright or inverted in separate runs to provide an additional control for low-level changes.

Since the base identity may become familiar after repeated presentations, the unfamiliar base identity from one run was never used as the new identity in a subsequent run for the same participant. Visual stimuli were projected on a screen at a spatial resolution of 1024 x 768 pixels and a frame rate of 60 Hz, at a viewing distance of 110 cm. They consisted of square images of 7.3° x 7.3° (14.1 x 14.1 cm) of visual angle. Participants viewed the images through a mirror placed on the head coil. They were instructed to stare at a small black fixation cross displayed at the center of the screen. While fixation and attention were not monitored, the large receptive fields of targeted face-selective regions (e.g., Kay et al., 2015) reduce the impact of small variations in fixation on the measured responses. The experiment was conducted using the EventIDE software (Okazolab).

### 2.4. Scanning sessions

Each participant underwent three scanning sessions separated by at least one week. In the first session, all participants performed 12 runs of the face categorization paradigm, with half of the runs displaying human faces and the other half monkey faces, alternating between the two categories. During the second and third sessions, participants completed conditions 1 and 2 of the face individuation paradigm, respectively. For each participant and each condition, 12 runs were acquired, including 6 runs with upright faces (3 with human faces and 3 with monkey faces) and 6 runs with inverted faces. Upright and inverted runs alternated, with two consecutive runs for each category (human or monkey faces). Each recording session lasted approximately 1 hour (i.e., 12 runs).

### 2.5. (f)MRI recordings

Recordings were performed on a 3 Tesla MR scanner (Philips Achieva) using a dedicated 32-channel head coil. At the beginning of each scanning session, anatomical volumes were acquired using a high-resolution T1-weighted magnetization-prepared gradient-echo image (MP-RAGE) sequence (repetition time, TR = 8.2 ms; echo time, TE = 3.7 ms; flip angle = 8°; FOV: 240 x 240 mm; voxel size = 1 x 1 x 1 mm; 180 sagittal slices). Functional images were collected with a T2*-weighted simultaneous multi-slice echoplanar imaging (EPI) sequence (TR = 1500 ms; TE = 30 ms; FA = 75°, SENSE factor = 1.6; multi-band, MB = 2; FOV = 240 × 240 mm; voxel size = 3 x 3 x 3 mm; matrix size = 80 × 80 mm; 48 slices acquired in an interleaved order).

### 2.6. Data processing

#### 2.6.1. Preprocessing of the anatomical and functional data

The volumes of each functional run were first motion-corrected with respect to the first volume of the first functional run of each session using trilinear interpolation. Across runs, the mean translational displacement was 3.6 ± 2 mm and the mean rotational movement was 4 ± 1°, no runs were excluded based on head motion. Since we used multi-band fMRI sequences, no slice-timing correction was applied. Individual anatomical data from the second and third sessions were co-registered with the anatomical images recorded in the first session, and all co-registered anatomical images from the three sessions were then averaged for each participant. Next, the motion-corrected functional data were co-registered to the individual average anatomical images. Functional and anatomical data were then aligned to the ACPC space using cubic spline interpolation and converted into Talairach space (Talairach and Tournoux, 1988). For each participant, an automatic segmentation based on the average anatomical images and an individual surface reconstruction were performed.

The volumetric functional data were finally projected onto the averaged individual cortical surface using a trilinear interpolation along the surface nodes (normal vector from −1 mm to +1 mm). All data preprocessing was performed using BrainVoyager software V23.0 (Brain Innovation, Inc., Maastricht, the Netherlands).

#### 2.6.2. Fourier analysis and amplitude of neural responses

For each condition, pre-processed fMRI data in the cortical surface space were first averaged across runs in MATLAB software (MathWorks^®^), using the Neuroelf toolbox to convert the data into a MATLAB-readable format. The corresponding 243 s time courses contained 27 cycles of our 9 s stimulus (see above). We removed the first cycle (9 s) to discard start-up transients (see e.g., Cottereau et al., 2012), leading to average time-courses of 234 s (i.e., 26 cycles of our stimulus). The averaged time-courses were subsequently smoothed using an iterative dilation process (n = 5) in which the value at each node was estimated from the average of the responses in its first-order neighborhood. This procedure corresponds to an effective surface smoothing of approximately 4 mm full-width at half-maximum (FWHM). Next, functional data were analyzed following the procedure described in Gao et al. (2018) and Laurent et al. (2023), by performing a Fourier analysis on the entire BOLD response time-courses and converting the amplitudes at the stimulation frequency (i.e., F = 0.111 Hz) and its second harmonic (2F = 0.222 Hz) into Z-scores, using the mean and standard deviation of the amplitude at 20 neighboring frequencies (10 on each side, with a bin width of 0.0043 Hz; Laurent et al., 2023). The Z-score was calculated by summing the baseline-subtracted amplitudes at the stimulation frequency and its second harmonic, relative to the standard deviation of the amplitudes at neighboring frequencies (Retter et al., 2021). This procedure was applied to the time course of each cortical node independently. A threshold of Z = 3.1 (*p* < 0.001, uncorrected) was selected as a conservative criterion, consistent with previous face perception studies (Kriegeskorte et al., 2007; Saygin et al., 2012; Gao et al., 2018; Gao et al., 2022).

#### 2.6.3. Defining activation and deactivation of neural responses

The activations and deactivations of the neural responses to the face categorization paradigm at the face stimulation frequency were defined by the phase of the BOLD response, as in Gao et al. (2018). For each individual, the histogram (20 bins) of phase values of all the nodes with a Z > 3.1 and with a positive phase value was calculated. The phase value of the histogram bin that has the largest number was used as the center phase (φ) to define all the nodes with their phase values within φ ± π/2 as activations (+ sign) and nodes with their phase values outside of this window as deactivations (−sign). A positive Z-score reflects nodes showing a significantly higher response to faces than to non-face stimuli (i.e., an increase in BOLD response amplitude), while a negative Z-score corresponds to nodes responding significantly less to faces than non-faces (i.e., a decrease in BOLD response amplitude). These signs were then applied to the thresholded Z-score maps to obtain the final response map. The final Z-score maps were displayed and visualized on individual cortical surfaces in BrainVoyager.

#### 2.6.4. Characterization of ROIs

For each participant’s brain, cortical regions of interest (ROIs) were defined based on BOLD responses to faces (i.e., regions showing greater activation for faces than for non-face objects) identified using our face categorization paradigm. Substantial interindividual variability has been widely observed in fMRI studies of face perception (Zhen et al., 2015; Gao et al., 2022), highlighting the importance of conducting analyses at an individual level. This independent functional localizer also helped prevent circularity (i.e., double-dipping) in subsequent analyses (Kriegeskorte et al., 2009). Hence, we bilaterally characterized the lateral MidFG (LatMidFG), corresponding to both mFus/FFA-2 and pFus/FFA-1; Weiner and Grill-Spector, 2010; Chen et al., 2023) the inferior occipital gyrus (IOG, corresponding to the OFA) the superior temporal sulcus (STS), the middle occipital gyrus (MOG), and the inferior frontal gyrus (IFG) in 10 participants. Table 1 summarizes the Talairach Coordinates of the peak node for each face-selective cluster and participant. We then computed the Z-scores for these regions from the face individuation paradigm. These Z-scores were obtained by averaging together the amplitude spectra of the local maxima of these areas and their fifth-order neighborhoods on the cortical tessellation (150 nodes were used on average).

**Table 1.** Talairach coordinates (x, y, z) of the peak node for each face-selective cluster identified in both hemispheres in 10 participants.

|  |  | mFus | pFus | IOG | pSTS | MOG | IFG |
|---|---|---|---|---|---|---|---|
| **S01** | **R** | 36 -39 -12 | 33 -58 -9 | 33 -71 -3 | 48 -44 19 | 38 -65 19 | 41 9 31 |
|  | **L** | -34 -40 -14 | -43 -64 -12 | -34 -87 -9 | -45 -51 9 | -39 -68 16 | -38 6 27 |
| **S02** | **R** | 36 -30 -16 | 35 -44 -13 | 30 -72 -7 | 50 -37 10 | 44 -56 11 | 42 13 28 |
|  | **L** | -36 -41 -15 | -44 -67 -12 | -30 -76 -5 | -57 -45 21 | -38 -54 9 | -33 1 25 |
| **S03** | **R** | 35 -43 -11 | 35 -65 -6 | 32 -70 -5 | 51 -36 12 | 48 -50 9 | 41 18 12 |
|  | **L** | -41 -39 -20 | -36 -66 -17 | -31 -82 -15 | -51 -38 -2 | -49 -70 9 | -42 22 33 |
| **S04** | **R** | 34 -50 -11 | 33 -55 -10 | 33 -68 -5 | 49 -40 5 | 39 -63 5 | 43 20 19 |
|  | **L** | -45 -49 18 | -39 -63 -15 | -40 -81 -15 | -44 -45 7 | -45 -71 12 | -36 16 28 |
| **S05** | **R** | 37 -32 -16 | 37 -55 -8 | 31 -75 -2 | 50 -41 3 | 43 -62 6 | 41 21 20 |
|  | **L** | -42 -57 -11 | -48 -69 -1 | -36 -76 -3 | -53 -42 19 | -33 -63 30 | -47 31 19 |
| **S06** | **R** | 34 -44 -12 | 33 -56 -5 | 35 -69 -6 | 47 -43 31 | 37 -67 12 | 40 19 10 |
|  | **L** | -41 -55 -13 | -35 -57 -13 | -37 -83 6 | -49 -34 10 | -44 -72 8 | -39 16 15 |
| **S07** | **R** | 38 -30 -21 | 39 -49 -20 | 40 -70 -12 | 49 -55 -1 | 46 -58 -5 | 36 30 34 |
|  | **L** | -38 -42 -20 | -36 -43 -19 | -42 -78 -8 | -44 -44 1 | -40 -60 -2 | -43 8 36 |
| **S08** | **R** | 34 -47 -12 | 30 -59 -8 | 30 -75 -2 | 50 -42 13 | 40 -66 15 | 45 10 15 |
|  | **L** | -34 -39 -19 | -38 -73 -14 | -34 -85 -5 | -41 -44 15 | -44 -80 3 | -45 7 28 |
| **S09** | **R** | 38 -39 -14 | 37 -50 -13 | 40 -81 -5 | 49 -31 15 | 52 -59 7 | 38 21 24 |
|  | **L** | -35 -49 -9 | -34 -51 -11 | -30 -75 -10 | -54 -48 12 | -42 -60 10 | -46 18 23 |
| **S10** | **R** | 38 -35 -12 | 36 -53 -7 | 33 -73 -2 | 47 -47 23 | 44 -58 13 | 40 23 25 |
|  | **L** | -37 -42 -16 | -39 -65 -8 | -35 -89 -5 | -48 -42 16 | -34 -73 25 | -44 23 26 |

#### 2.6.5. Statistical analysis

We compared the Z-scores obtained in all face-selective clusters during upright runs to those obtained during inverted runs. The significance of the inversion effect was assessed using a permutation test. For each cluster, the difference conditions (upright vs. inverted) were randomly shuffled at the subject level 10000 times, with the condition labels randomly reassigned for each subject in each permutation, and the signed Z-score differences were recomputed for each permutation. This procedure generated a normal distribution of Z-score differences with an equal probability of positive and negative differences in the absence of any effect. Observed differences exceeding the 95th percentile of this distribution were considered reliably significant.

## 3. Results

### 3.1. The cortical face-selective network revealed by frequency-tagging

The frequency-tagging face categorization paradigm aimed to localize individual face-selective regions in all participants. Figure 2A shows the face-selective activations and deactivations projected on inflated reconstructions of the left and right cortical hemispheres of two representative participants, S02 and S03 (see Supplementary Figure 3 for the face-selective regions defined in all individuals). These fMRI activations were estimated from all runs, combining human and monkey face stimuli. Clusters with robust face-selective responses (Z > 3.1, *p* < 0.001, uncorrected) were observed in typical regions previously described in neuroimaging studies (Grill-Spector et al., 2004; Weiner and Grill-Spector, 2010; Zhen et al., 2015; Gao et al., 2018). Hence, we were able to objectively define the ventral core face-selective network in all participants’ VOTC, consisting of the OFA in the IOG (Pitcher et al., 2011) and two FFAs in the MidFG (mFus/FFA-2, located more anteriorly, and pFus/FFA-1, located more posteriorly, according to the terminology of Weiner and Grill-Spector, 2010; Weiner et al., 2014; Grill-Spector et al., 2017). Yet, given that fMRI activations appear to be continuous along the fusiform gyrus in some individuals (e.g., in S04, S06, and S07; Supplementary Figure 3), we increased the statistical threshold to define the two local maxima along the MidFG in all participants. Nevertheless, the most significant response at the face-selective frequency (i.e., 0.111 Hz; see the amplitude spectra in Figure 2B) was objectively quantified in either mFus/FFA-2 or pFus/FFA-1 in 18/20 hemispheres (only participants S02 and S05 showed stronger IOG responses in only one of their hemispheres). These activations were associated with typical Talairach coordinates of FFA-1 and FFA-2 (38, −44, −15 and 37, 58, −12 respectively, compared, for instance, to the coordinates reported in Pinsk et al., 2009: 40, −43, −14 and 39, −57 −13). Furthermore, in all individual brains, we observed that fMRI activations were consistently located in the lateral portion of the MidFG, while fMRI deactivations fell in the medial part of the MidFG. This observation is in line with the anatomical and cytoarchitectonic boundaries of face-selective activations defined by the MidFG (Weiner et al., 2017; Weiner, 2019).

**Figure 2.**
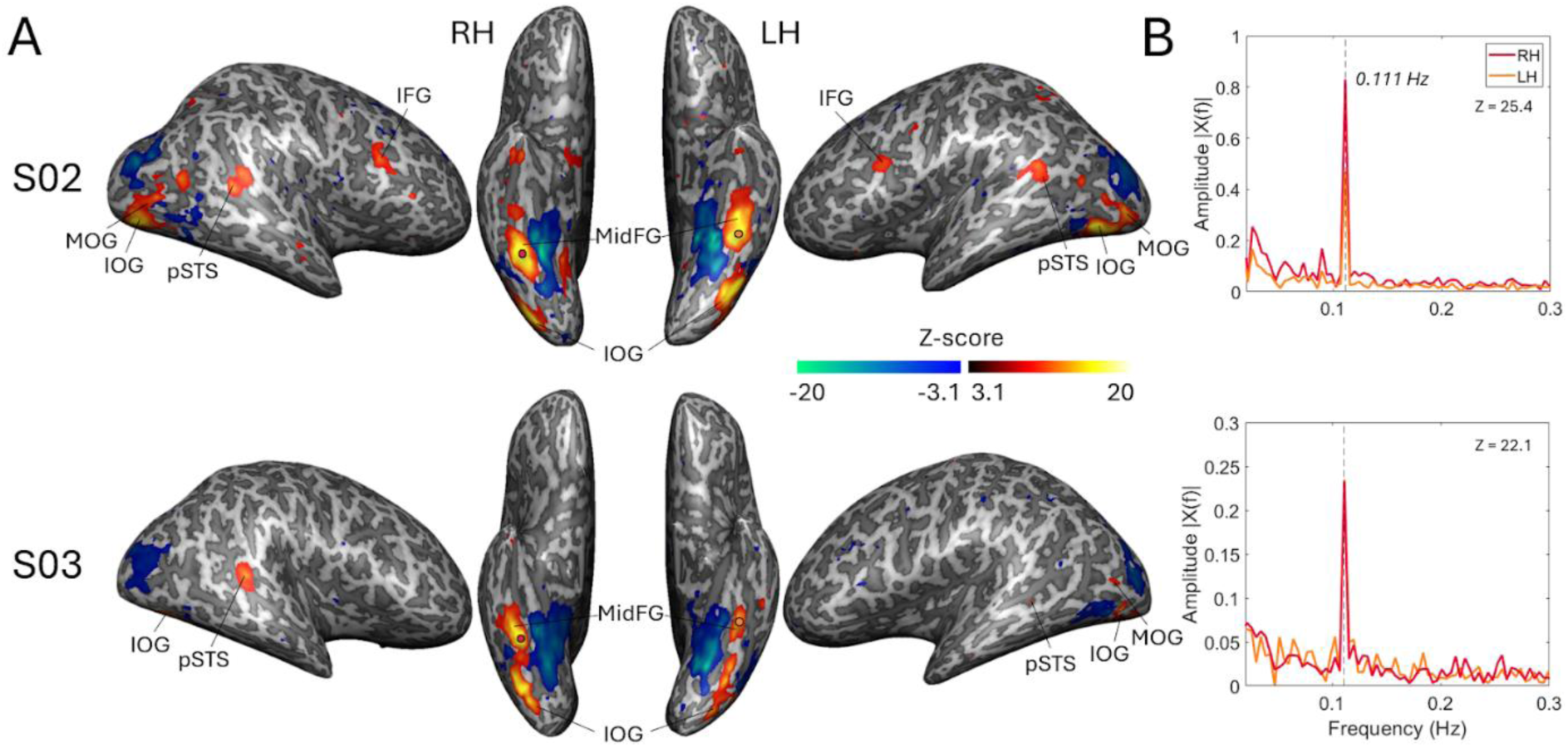
Face-selective clusters identified by the frequency-tagging face categorization paradigm in two representative participants. **(A)** Significant face-selective activations (hot colors) and deactivations (cold colors) projected onto lateral and ventral views of the inflated cortical hemispheres of S02 (top panel) and S03 (bottom panel). Data were thresholded at Z = 3.1 (i.e., *p* < 0.001, uncorrected). LH: left hemisphere, RH: right hemisphere. **(B)** The associated amplitude spectra of the BOLD responses around the local maxima of MidFG (represented by dots in panel A) are shown in the rightward panels for the right (in red) and left (in orange) hemispheres. Note the high SNR of the BOLD activity at 0.111 Hz (indicated by the gray dotted line). The averaged Z-scores at the local maxima across both hemispheres are provided in the upper-right corner of each graph.

Besides the ventral face-selective regions, we also identified significantly higher activations to faces than non-face objects in areas commonly seen in typical adults, such as the pSTS (Haxby et al., 2000; Duchaine and Yovel, 2015), IFG (Fox et al., 2009; Ishai et al., 2005), and MOG. Despite a relative degree of consistency across most participants, not all of these areas were identifiable in both hemispheres of the 10 participants (see Section 2.6.4.), and substantial variability in ROI size was observed across individuals. This finding is consistent with the genuine interindividual variability described in the number and size of face-selective regions in humans (Zhen et al., 2015; Gao et al., 2022). At a more liberal threshold (Z > 1.64, *p* < 0.05, uncorrected), we identified an additional bilateral cluster in the ventral anterior temporal lobe (vATL) of only one participant (i.e., S02). The lower signal-to-noise ratio (SNR) observed in this region is likely due to magnetic susceptibility artifacts in the vATL, arising from the proximity of the auditory canals (Wandell, 2011; Axelrod and Yovel, 2013). Given the absence of fMRI-defined vATL in a majority of participants and the lack of identity responses identified with this paradigm (Supplementary Figures 4 and 5), further analyses did not focus on this region as an ROI. Otherwise, all face-selective regions were characterized by a very high SNR (e.g., Z-scores ranged from 4.2 to 53.5, mean ± standard deviation = 18.6 ± 11.8 at the peak face-selective node within the LatMidFG). At the group level, we observed the strongest response in pFus/FFA-1 (mean ± standard deviation = 19.7 ± 13.1 at the peak face-selective node).

Our face categorization paradigm successfully identified the highest SNR in the right LatMidFG of these two representative participants, consistent with previous neuroimaging studies (Sergent et al., 1992; Kanwisher et al., 1997; Rossion et al., 2012; Gao et al., 2018, 2022). In these representative participants, amplitude spectra derived from a Fourier transform of the BOLD responses revealed higher SNR at the face-selective frequency in the right mFus/FFA-2 compared with the left. However, among all participants, only half showed a right hemispheric dominance (i.e., S02, S03, S04, S07, and S10; Supplementary Figure 3) whereas face-selective activations in the remaining participants were more bilaterally distributed. This finding aligns with substantial individual variation in the lateralization of the face perception network (Thome et al., 2022).

### 3.2. Responses to unspecific changes in face identity in the VOTC

In our first condition (i.e., same image of the base identity presented), robust unspecific (FIR) responses were identified across all face-selective ROIs previously defined from our face categorization paradigm and across all participants. Although activations were relatively small in participant S05 (see Supplementary Figure 4 for all individual responses), the lowest SNR was also observed in this subject for the face categorization paradigm, suggesting reduced signal in this individual. Figure 3A presents the corresponding activations on inflated reconstructions (estimated from all the runs with human and monkey faces combined) of the left and right cortical hemispheres of S02 and S03 (all Zs > 5, *p* < 10^−6^, uncorrected). As illustrated in Figure 3B for two representative subjects, high SNR was observed at the identity-change frequency (i.e., 0.111 Hz) in the LatMidFG of all participants except S05. We observed the strongest unspecific face individuation responses in the pFus/FFA-1 (Z-scores ranged from 1 to 32.2, mean ± standard deviation = 16.1 ± 3.1, averaged across hemispheres) and the mFus/FFA-2 (Z-scores ranged from 1 to 19.9, mean ± standard deviation = 12.7 ± 2, averaged across hemispheres).

**Figure 3.**
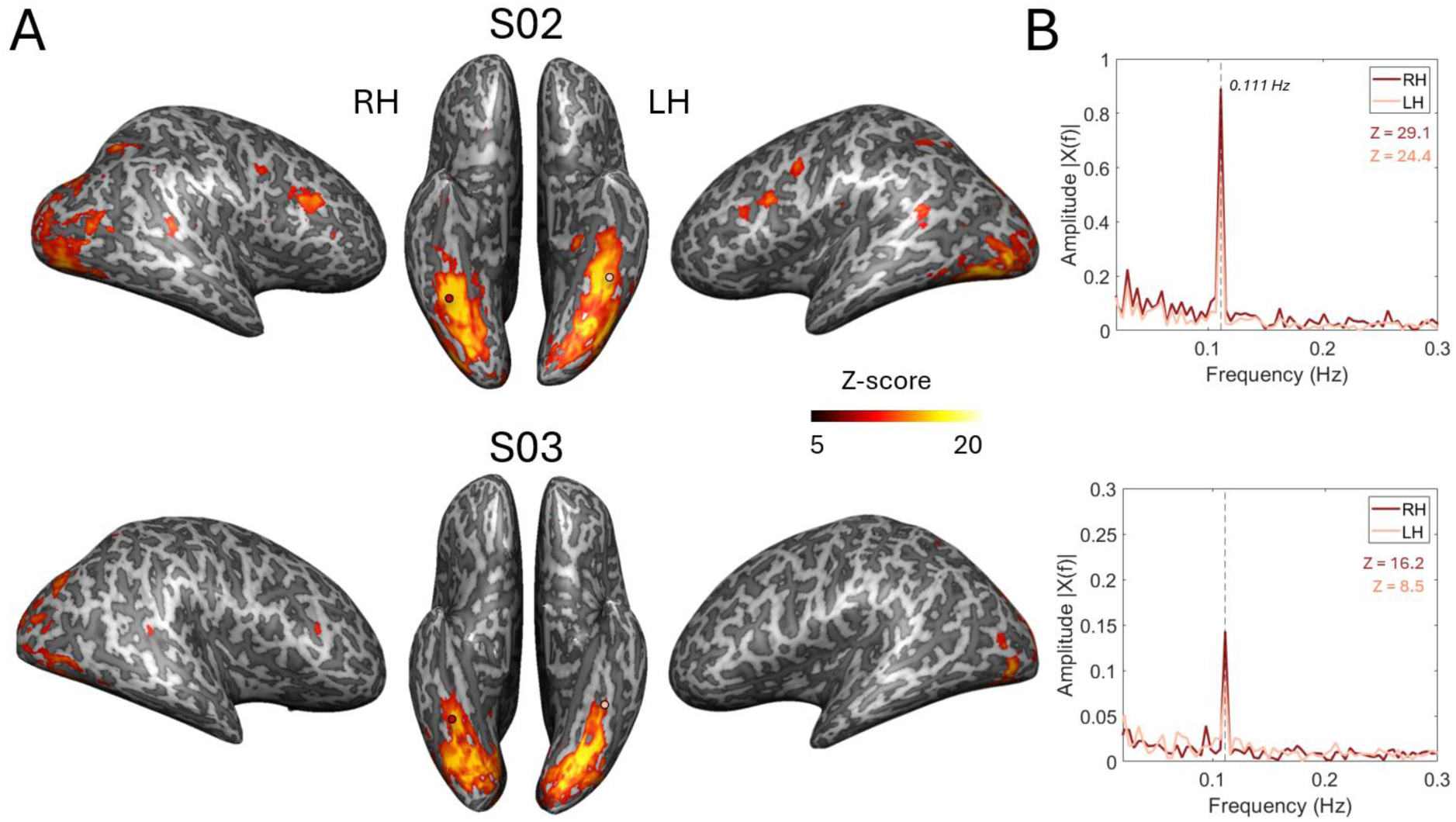
Unspecific face individuation responses across low-level changes in two representative participants. **(A)** Significant face-selective activations projected on the inflated cortical hemispheres of S02 (top panel) and S03 (bottom panel) for unfamiliar faces (all Zs > 5). LH: left hemisphere, RH: right hemisphere. **(B)** The associated amplitude spectra of the BOLD responses around the local maxima of the LatMidFG identified from the face localizer (indicated by dots) are shown on the rightward panels for the right (dark pink) and left (light pink) hemispheres. The averaged Z-scores in both hemispheres for upright faces are provided in the upper-right corner of each graph.

Hence, discriminating new and different identities elicited significant fMRI activations at the exact frequency of identity changes, with the largest responses located in the LatMidFG and IOG, corresponding to FFA-2, FFA-1, and the OFA, respectively. This observation is consistent with a previous neuroimaging study using a similar approach (Gao et al., 2019), and leading to large responses selective to identity changes in the occipito-temporal cortex on the scalp, as recorded in prior EEG studies (Liu-Shang et al., 2014; see Rossion et al., 2014 for review). In addition, in the two participants depicted in Figure 3, the highest SNR among the LatMidFG ROIs was observed in the right hemisphere. However, the whole-brain maps did not show a consistent hemispheric difference in face-selective responses, and no significant hemisphere lateralization was found at the group level.

### 3.3. Face identity responses preserved across changes in viewpoints

Next, we examined whether these unspecific responses related to identity changes can generalize to higher-level changes of the same base identity occurring throughout the cycle (i.e., background, size, head position and orientation, gaze direction, etc.). As in the first condition, FIR responses were observed in core face-selective ventral regions (FFA-1, FFA-2), although these were less extensive and more clustered around the peak node for this condition (e.g., Z-scores ranged from 0.6 to 12.6, mean ± standard deviation = 5.9 ± 1.4 in pFus/FFA-1, Z-scores ranged from 1.6 to 11.8, mean ± standard deviation = 5.3 ± 1.1 in mFus/FFA-2, averaged across hemispheres). Figure 4 presents the respective activations on inflated cortical surfaces of the left and right cortical hemispheres of S02 and S03 (all Zs > 3.1, *p* < 0.001, uncorrected) and the amplitude spectra associated with the LatMidFG. When human and monkey faces runs were analyzed independently, the same cortical areas were activated across low-and higher-level face identity changes, although the network associated with monkey faces appeared more spatially extended in some participants (Supplementary Figures 6 and 7). Therefore, all runs were combined in subsequent analyses to increase the global SNR, as further addressed in the Discussion section. For both participants, the peak in this face-selective region was higher in the right than in the left hemisphere, although the whole-brain maps did not show a consistent hemispheric difference in face-selective responses. At the individual level, all participants except S05 (for the same reasons as discussed in Section 3.2.) exhibited strong face individuation responses (see Supplementary Figure 5 for all individual responses), primarily localized in the LatMidFG and IOG. Importantly, we also found specific responses to face identity recognition in the IFG of 6/10 participants (in at least one hemisphere, e.g., S02 in Figure 4). These results demonstrate generalization of the FIR response across changes in viewpoint in these specific face-selective regions.

**Figure 4.**
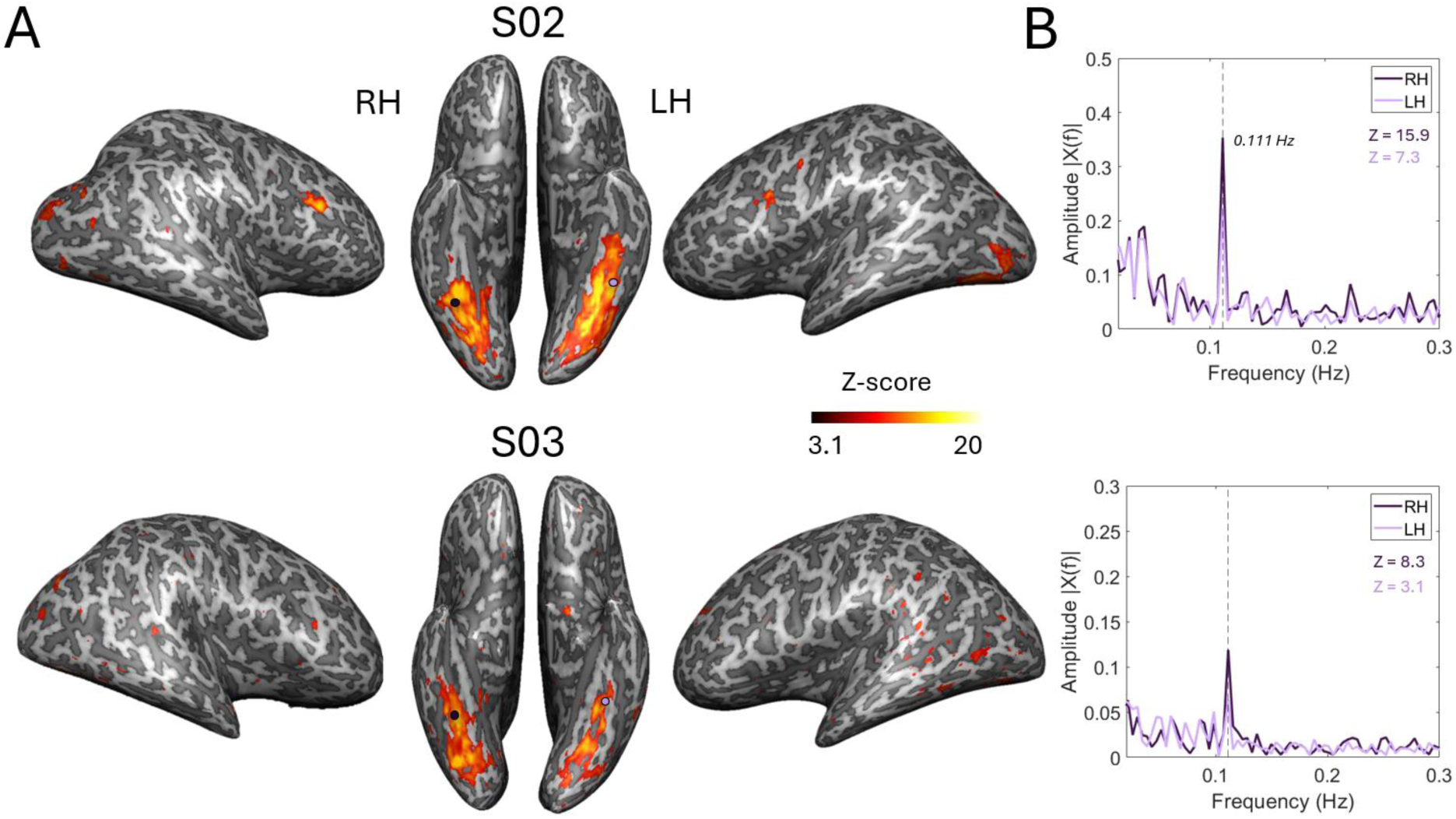
Generalized face individuation responses in two representative participants. **(A)** Significant face-selective activations projected on the inflated cortical hemispheres of S02 (top panel) and S03 (bottom panel) for unfamiliar faces (all Zs > 3). LH: left hemisphere, RH: right hemisphere. **(B)** The associated amplitude spectra of the BOLD responses around the local maxima of LatMidFG from the face localizer (indicated by dots) are shown on the rightward panels for the right (dark purple) and left (light purple) hemispheres. The averaged Z-scores in both hemispheres for upright faces are provided in the upper-right corner of each graph.

### 3.4. High-level processing during unfamiliar face individuation

Although FIR responses varied substantially across individuals, robust image-based face-discrimination responses were observed at the group level across both conditions in the core face-selective ventral regions (Figure 5; see Supplementary Figure 8 for separate hemispheres). We therefore quantified the face individuation response for each region, with a maximal response located in the LatMidFG, specifically within pFus/FFA-1. While the response SNR decreased under image variations, responses remained largely significant in the LatMidFG (mFus and pFus) and IOG (Z > 3.1, *p* < 0.001, uncorrected). The same pattern of fMRI activations was observed using a Bonferroni-corrected threshold at *p* < 0.05 across the six ROIs (corrected α = 0.008, corresponding to Z > 2.39, corrected), demonstrating the robustness of these results. In contrast, responses in the pSTS and low-level visual regions such as the MOG or V1 (Supplementary Figure 9) were negligible in the second condition. This finding was consistent across all participants and was observed for both human and monkey faces (Supplementary Figure 11). A repeated-measures ANOVA revealed a significant ROI x Condition interaction (*F*(5, 45) = 4.42, *p* = 0.002, ηp² = .329), indicating that the effect of condition varied across regions. Consequently, this condition provides a measure of generalization of the FIR to unfamiliar faces, as the response elicited by each picture of a novel facial identity must consistently differ from that elicited by the repeated identity presented under varying viewing conditions. This result is consistent with previous EEG studies demonstrating viewpoint-invariant neural representation during face individuation response for unfamiliar faces (Damon et al., 2020; Yan et al., 2020).

**Figure 5.**
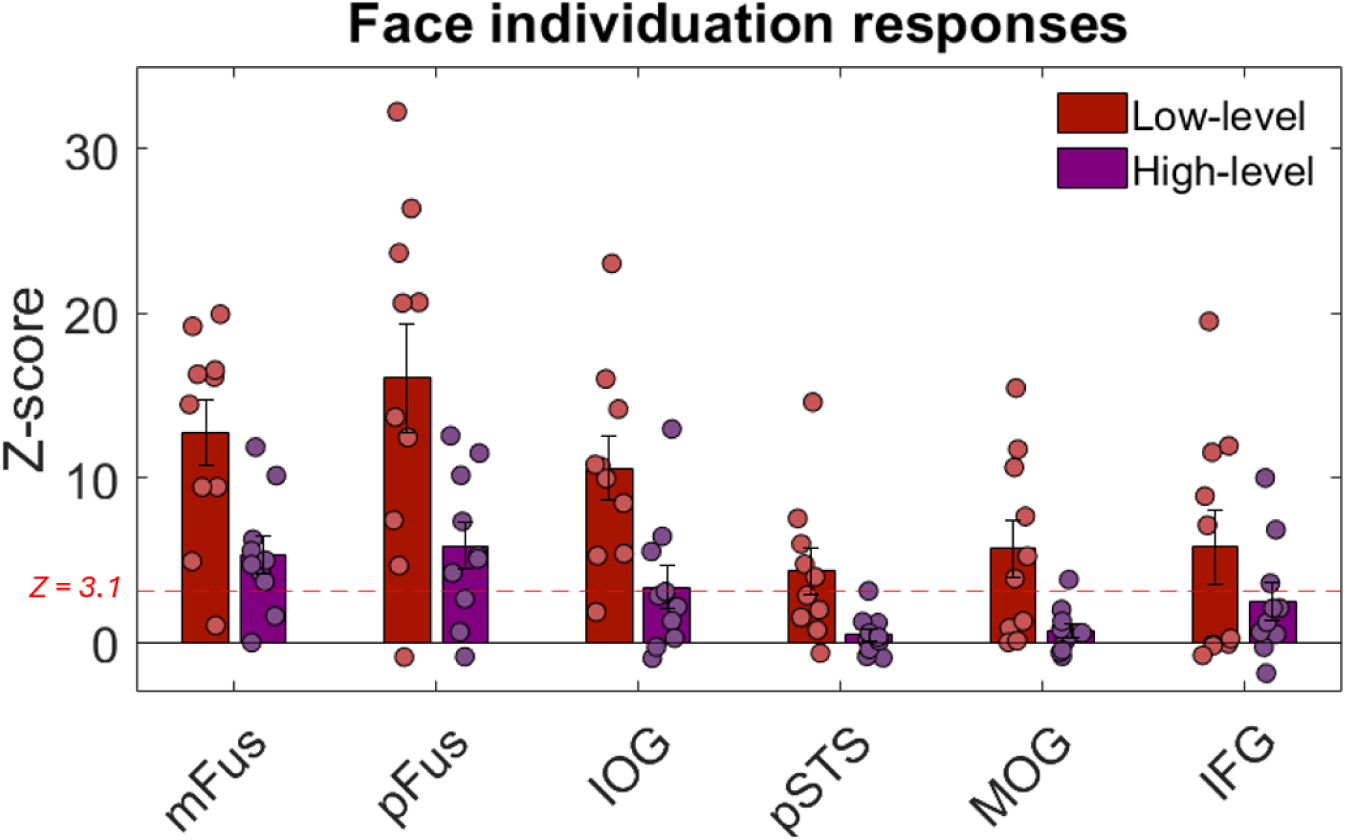
Unspecific and generalized face individuation responses to unfamiliar faces. Grouped bars show the mean Z-scores for each face-selective cluster in response to identity changes across low-level (red) and high-level (purple) conditions. Data was averaged across left and right hemispheres. Error bars represent the standard error of the mean. The horizontal dotted line represents a threshold level of Z > 3.1, *p* < 0.001, uncorrected; *p* < 0.01 with Bonferroni correction.

Face individuation responses for inverted faces were still observed with a high SNR at the stimulation frequency in the core ventral face-selective regions (Supplementary Figures 4 and 5). However, fMRI activations reflecting individual face discrimination in the LatMidFG were generally larger and more consistent for upright faces (Figure 6; see Supplementary Figure 10 for separate hemispheres). The inversion effect in mFus was significant in both conditions based on permutation testing and was largely invariant to high-level stimulus changes (*p* = 0.028, 95% CI [0.8-4.9] for low-level changes; *p* = 0.008, 95% CI [0.9-3] for higher-level changes). To account for multiple comparisons, Bonferroni correction was applied across the six ROIs (corrected α = 0.00833), the inversion effect remained significant for higher-level changes, supporting the reliability of this effect. In particular, some participants showed a substantial reduction in responses to inverted faces across high-level identity changes (e.g., S03 and S09; Supplementary Figure 5). Interestingly, face discrimination responses in the IFG also tended to be reduced for inverted faces during low-level changes. Half of the participants showed a significant decrease in BOLD response for the first condition, even though the effect did not reach significance at the group level (permutation test, *p* = 0.076). This trend did not persist across high-level image variations. In the IOG, no reliable difference was found between upright and inverted face identities (*p* = 0.553). Overall, only 2 participants showed significantly reduced activations for inverted faces in this face-selective region. Face individuation responses in the IOG thus appear to be largely unaffected by picture-plane inversion.

**Figure 6.**
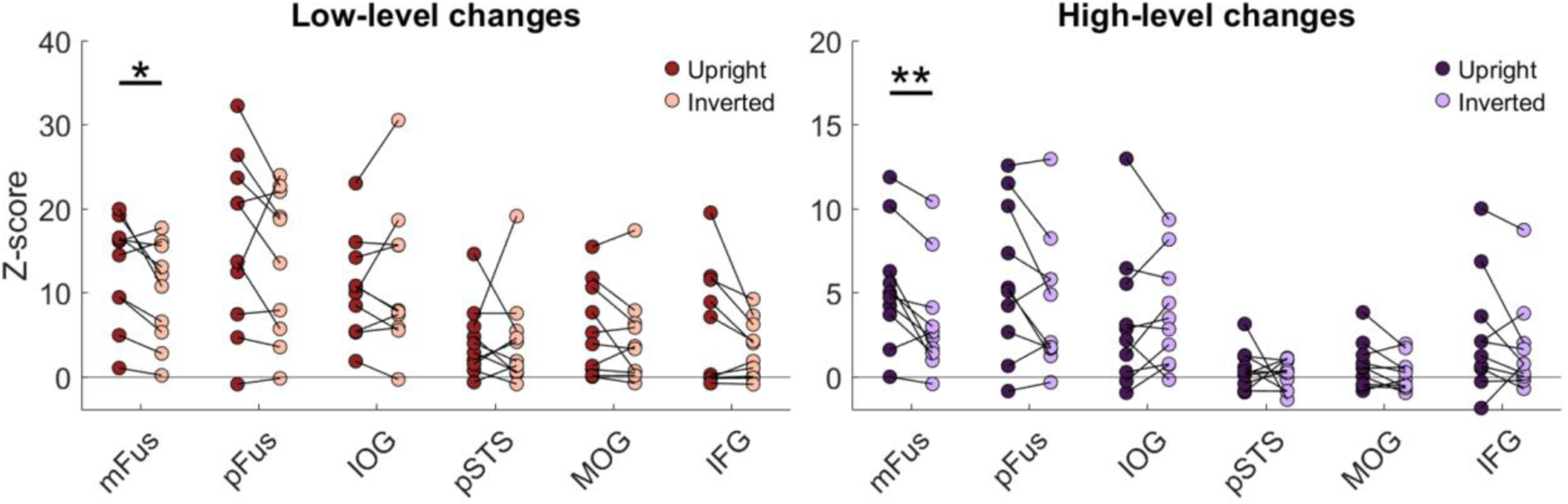
Unspecific and generalized face individuation responses to upright and inverted unfamiliar faces. For each subject, colored dots provide the mean Z-scores for each face-selective cluster in response to identity changes, with FIR responses across low-level changes on the left panel (red: upright faces; light red: inverted faces) and high-level changes on the right panel (purple: upright faces; light purple: inverted faces). Data was averaged across left and right hemispheres. \**p* < 0.05; \*\**p* < 0.01.

To further characterize face individuation responses to human and monkey faces separately, we analyzed responses to upright and inverted faces for each stimulus set (Supplementary Figure 11). FIR responses to low-level changes were observed in all core face-selective regions for both human and monkey faces. In the high-level changes condition, significant FIR responses were also retained in the mFus and pFus for both stimulus sets (*p* < 0.05, uncorrected). However, permutation tests revealed no significant inversion effect in any ROI for either human or monkey faces, likely due to the reduced SNR resulting from analyzing the datasets separately. Consistent with the individual whole-brain maps, responses to monkey faces were significantly larger than those to human faces in the pFus and IOG (permutation tests, *p* < 0.01), a difference that may be due to several factors such as the fact that monkey faces presented across identity variations (ID1 to ID12; Supplementary Figure 2) and alternates came from different image sets (with potentially female faces in the alternate category) but also greater within-identity image variability for human faces in terms of viewing conditions (lighting, texture, background, etc.), and other image properties (Figure 1).

## 4. Discussion

### 4.1. The value of fMRI frequency-tagging for measuring face individuation

Face Identity Recognition (FIR) is typically conceived as the ability to identify a familiar face and its corresponding specific semantic information. Since images of unfamiliar faces are more difficult to match by identity than images of familiar faces, unfamiliar FIR has sometimes been considered unnatural and primarily driven by low-level cues (Hancock et al., 2000; Megreya and Burton, 2006; Jenkins et al., 2011; Young and Burton, 2018). However, successful FIR fundamentally depends on the capacity to visually individuate faces, including unfamiliar ones (Rossion, 2018). At the neural level, although numerous neuroimaging studies have investigated the neural basis of FIR with pictures of unfamiliar faces, the effects found, notably with MVPA-fMRI, often reflect image-based discrimination within restricted sets of stimuli, and therefore do not provide clear evidence for identity-level generalization of unfamiliar faces.

Here we measured unfamiliar face individuation with a fMRI frequency-tagging approach that provides highly sensitive, objective and reliable face-selective responses (Gao et al., 2022; Laurent et al., 2023). Although conceptually related to fMRI adaptation paradigms, this approach offers important methodological advantages, including objective quantification of identity-change responses at a pre-defined frequency, without requiring BOLD response modeling (Gao et al., 2018). Its primary contribution therefore lies in providing a sensitive and objective method for measuring face individuation. The present study relies on univariate analysis, but the paradigm also allows for multivariate approaches, which could provide complementary information about identity representations (but see Rossion, 2014; Dubois et al., 2015). Using this model-free analysis in the Fourier domain, we found robust (i.e., high SNR) image-based FIR responses localized bilaterally in core face-selective ventral regions, the lateral MidFG (corresponding to FFA-1 and FFA-2), and the IOG (including the OFA), across all participants. Importantly, these responses were identified and quantified at the identity-change frequency (i.e., 0.111 Hz), enabling objective measurement of neural activity associated with visual face individuation, consistent with prior work on face categorization (Gao et al., 2018, 2022; see also Laurent et al., 2023 for a study in non-human primates). Furthermore, the optimal rapid presentation rate (i.e., 6 Hz; Alonso-Prieto et al., 2013; Gentile & Rossion, 2014; Retter et al., 2021), offers several advantages. First, it allows only a single gaze fixation per image due to the fixation cross constraint. Second, combined with the presentation of a large number of natural individual face identities and comparison with inverted-face stimulation, it provides controls for the contribution of low-level cues.

Significant FIR neural responses were observed in the LatMidFG and IOG, in agreement with the regions typically identified in previous fMRI adaptation studies (Gauthier et al., 2000; Grill-Spector and Malach, 2001; Davies-Thompson et al., 2009; Gentile and Rossion, 2014). Only one participant (S05) showed a relatively low overall SNR, which was consistent across all brain regions and conditions. We also found robust responses to identity changes in the IFG in the low-level condition, although these responses were not reliably maintained with high-level variations. While these latter results are consistent with prior fMRI studies using famous faces (Leveroni et al., 2000; Ishai et al., 2005), they provide additional evidence for the recruitment of prefrontal regions in the discrimination of newly encountered faces. Interestingly, cortical networks were highly similar when analyzing human and monkey faces independently, with the same cortical areas being activated (Supplementary Figure 4). We also reported variability in spatial extent, anatomical location, and the number of clusters involved in identity discrimination across participants (Figure 3). Differences in individual expertise for face individuation or fluctuations in SNR may contribute to this variability. Importantly, such inter-individual variability may also provide insight into genuine anatomo-functional differences underlying face recognition across individuals (Gao et al., 2022).

### 4.2. Discrimination and generalization supporting unfamiliar face individuation

The present study goes beyond previous work by including a key condition that introduces stimulus changes either at low-level (e.g., background) or at high-level (i.e., head orientation, gaze direction, etc.). While activations to identity changes without such variations were observed also in early visual cortex and in the pSTS, these responses did not resist to low- and high-level stimulus variations (Figures 4 and 5). The absence of reliable responses in the pSTS in this condition indicates that head orientation variations are neutralized and that presentation of faces from different viewpoints is not interpreted as motion of the individuals, probably due to substantial size and position variation of the face in our images. The reduction in responses may also have been influenced by differences in several stimulus properties between the two conditions, including the variability of low-level visual cues and head and gaze orientation. In contrast, robust responses were still observed in the ventral areas of the LatMidFG and IOG, supporting viewpoint-generalized responses in these regions. Indeed, in condition 2 (‘generalized’), a FIR response can only emerge significantly if the system detects identity changes generalized across viewpoint variations of the same identity (Yan et al., 2020 for a study using familiar faces). Since the faces presented here were unfamiliar identities, long-term familiarization may further enhance these generalized FIR responses, increasing response magnitude and recruiting an extended cortical network (Zheng et al., 2026). Although repeated exposure to the base identity may facilitate the detection of novel identities through short-term familiarization, no systematic increase in BOLD response amplitude was observed at the group level (Supplementary Figures 12 and 13). Accordingly, this second condition specifically isolates more ecologically valid neural FIR responses, while demonstrating generalization across viewpoint changes in core face-selective ventral regions (Figure 5). The observed responses are consistent with discrimination between different individual faces, although they may also partly reflect natural variations between face images that cannot be dissociated from identity.

A limitation of the present design is that the low-level and high-level conditions were tested in separate scanning sessions and were not counterbalanced. Differences in response magnitude between the two conditions may therefore also be influenced by session-related factors. Despite this, reliable identity-generalized responses were detected in the generalized condition. Another limitation relates to the use of natural face images, which contain a wide range of within-identity and between-identity variations (Supplementary Figures 1 and 2). These include differences in age and gender, as well as variations in lighting, texture, hairstyle, or facial hair. Because the images were obtained from publicly available image sources, these characteristics could not always be systematically controlled. Such variability may be particularly pronounced in human faces, which exhibit greater diversity in appearance and context. To account for this variability, we used a large and diverse set of images and included an inverted-face condition to control for the contribution of low-level image variations to the observed responses.

Separate analyses of responses to human and monkey faces confirmed that both stimulus sets elicited strong activations within the core face-selective network, although stronger identity-related responses for monkey faces were observed. Since faces of genetically homogeneous populations are much more likely to differ in humans than other animal species (Sheehan & Nachman, 2014) and humans lack expertise in distinguishing monkey faces, this difference may be due to two factors. First, a larger between-identity variability in our non-human primates stimulus set. Second, human faces may also exhibit greater within-identity variability in appearance and context (e.g., hairstyle, facial hair, or skin tone), which could make identity discrimination more challenging.

Importantly, we also observed no reliable responses in the vATL and the temporal pole. Although fMRI has known limitations in exploring anterior regions beyond the MidFG due to magnetic susceptibility artifacts (Rossion et al., 2024), these limitations may contribute to the absence of FIR responses in these regions. However, the lack of responses in the most anterior regions could also reflect the use of unfamiliar faces lacking associated semantic information, typically involved in temporal lobe activations (Collins and Olson, 2014; Rice et al., 2018). Overall, our findings are in line with previous studies using similar paradigms in EEG (Liu-Shang et al., 2014), intracerebral recordings (Jacques et al., 2020), and fMRI (Gao et al., 2019), and provide strong evidence for high-level discrimination of unfamiliar faces in the human VOTC.

### 4.3. Unfamiliar face identity recognition relies on high-level neural processes

We further examined the high-level neural processes underlying FIR by exploiting the effect of picture-plane inversion. Given that human behavioral performance at FIR is significantly impaired for inverted faces (Yin, 1969; Freire et al., 2000; Rossion, 2008, for review), this manipulation allows us to isolate processes that are not limited to the discrimination of physical differences between individual faces, preserved under inversion. At the individual level, 80% of participants showed a significantly reduced FIR response in the LatMidFG (either in mFus or pFus) for inverted stimuli (both human and monkey faces) without controlling for low-level and high-level variations not related to identity, and 70% exhibited a reliable reduction that persisted under high-level changes (Figure 6). Consistent with this, fMRI responses associated with face individuation were generally stronger in the MidFG, particularly in mFus, which showed a significant inversion effect at the group level. The inversion effect was no longer observed when responses to human and monkey faces were analyzed separately, due to the reduced SNR in these analyses. These findings directly confirm that the observed responses cannot be attributed solely to objective physical differences and challenge the view that unfamiliar faces are individuated based on low-level cues. Nevertheless, robust FIR responses were still observed for inverted faces in the core ventral face-selective regions and the IFG, indicating that face individuation mechanisms in these regions remain engaged even when faces are presented upside-down (Yovel and Kanwisher, 2005; Mazard et al., 2006; Gilaie-Dotan et al., 2010; James et al., 2013). In this context, the face inversion effect as measured with fMRI appears to reflect enhanced persistence of neural responses (Strother et al., 2011), along with altered dynamics of recovery from adaptation (Mazard et al., 2006).

## 5. Conclusions

Overall, our findings illustrate the validity and efficiency of fMRI frequency-tagging approach for characterizing the cortical network underlying unfamiliar face recognition. Image-based FIR responses were objectively identified in core face-selective ventral regions, in agreement with previous fMRI studies. By including a condition requiring identity discrimination and generalization across image variations, we found evidence that viewpoint-generalized responses are supported by ventral regions of the cortical face network. The persistence of identity-changes responses, together with a reliable inversion effect in the LatMidFG, argues against a purely low-level cue-based account of unfamiliar face individuation. This functional paradigm could be extended to directly compare unfamiliar FIR mechanisms in human and non-human primates (with the frequency-tagging approach already validated in monkeys; Laurent et al., 2023) and further explore the neural basis of familiar FIR.

## Ethics statement

All subjects gave written informed consent to participate to the study and the protocol was approved by the French Ethics Committee of the CPP Sud-Ouest Outre-Mer IV (N°2015-A01801-48).

## Data and Code Availability

FMRI data will be provided online (Open Science Framework) upon acceptance of the paper (http://xxx.xxx). Analysis codes are accessible at the following link: https://osf.io/6r8wz/. The stimuli are also available online: https://osf.io/ds6zm/overview?view_only=c6c1a741f3f94793b9dce6f463708ac4.

## Authors Contributions

M.-A.L., X.G., B.R., and B.R.C. designed research; M.-A.L. and B.R.C. performed research; M.-A.L. analyzed data; J.-B.D. contributed analytic tools; N.V. and I.B. provided resources; M.-A.L., B.R., and B.R.C. wrote the original manuscript; M.-A.L., J.-B.D., N.V., I.B., X.G., B.R., and B.R.C. reviewed and edited the manuscript; B.R.C. and B.R. acquired funding.

## Acknowledgments

We sincerely thank the participants for their involvement in the study.

