## Supplementary Material for "The neural basis of unfamiliar face recognition in humans with fMRI frequency-tagging"

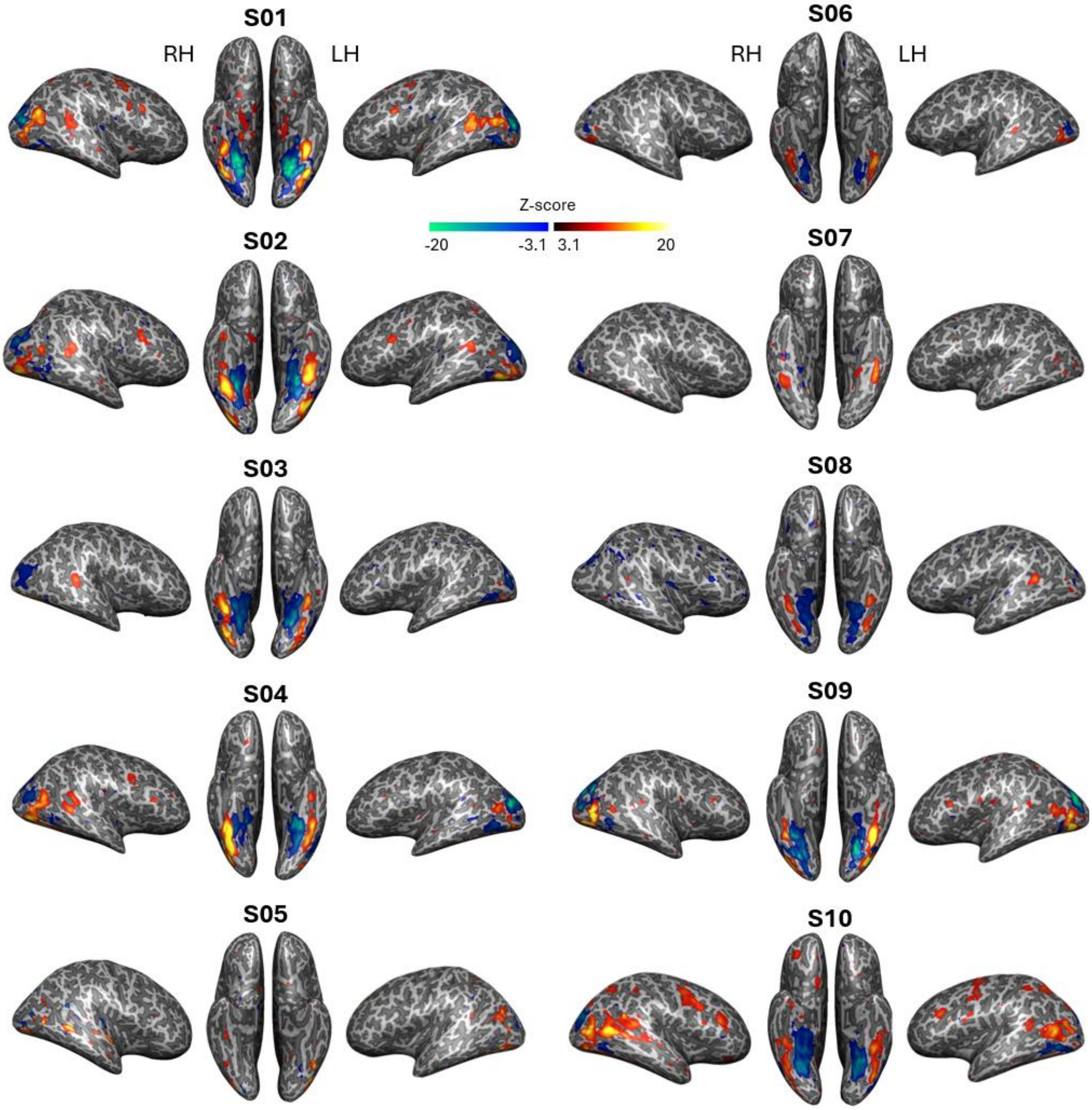


**Supplementary Figure 3. Face-selective clusters identified by the frequency-tagging face categorization paradigm in all participants.**
Significant face-selective activations (hot colors) and deactivations (cold colors) projected on the inflated cortical hemispheres of all participants. Lateral and ventral views of brains are shown here. Data were thresholded at Z = 3. LH: left hemisphere, RH: right hemisphere.

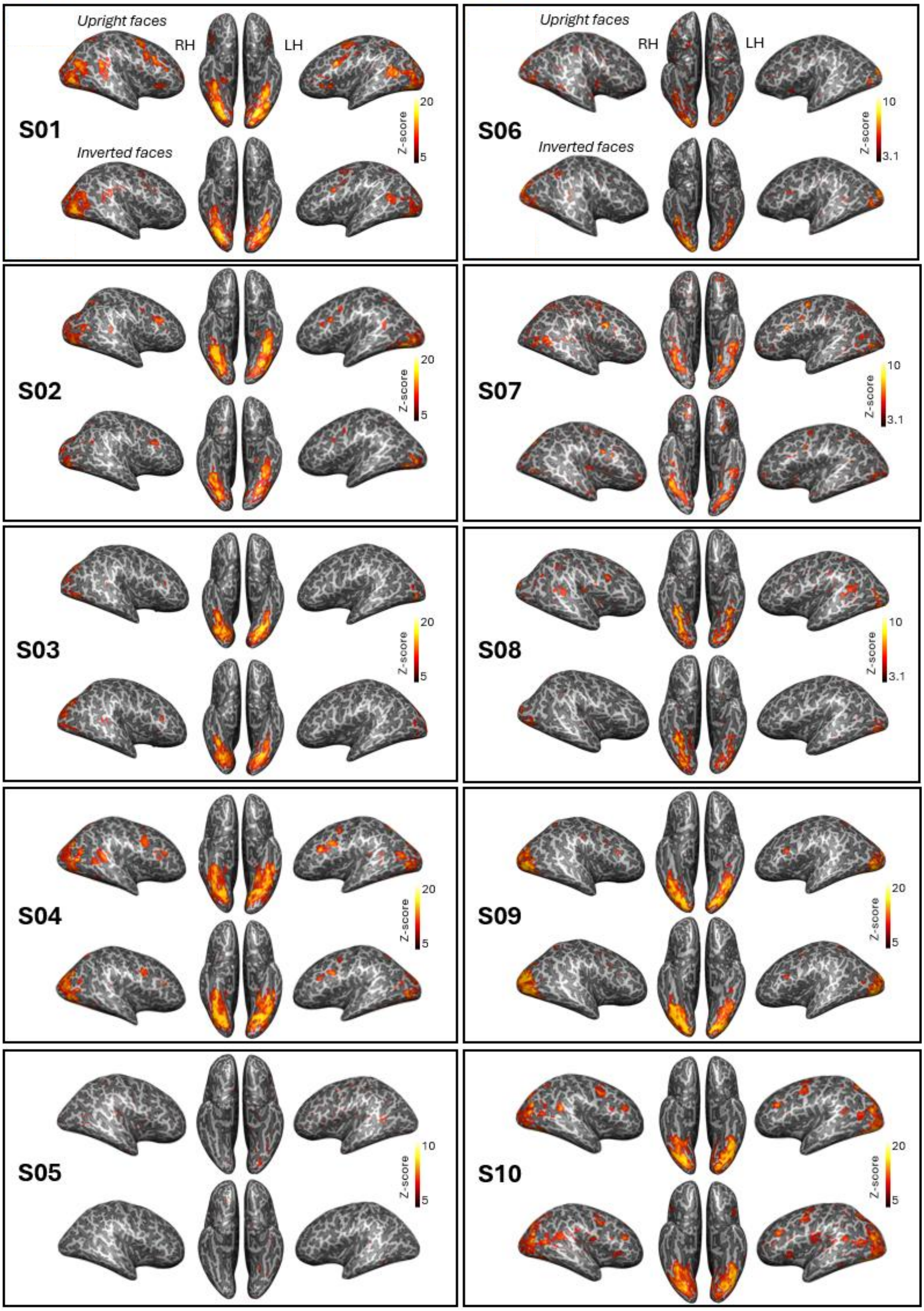


**Supplementary Figure 4. Face individuation responses for upright and inverted faces across low-level changes.** LH: left hemisphere, RH: right hemisphere.

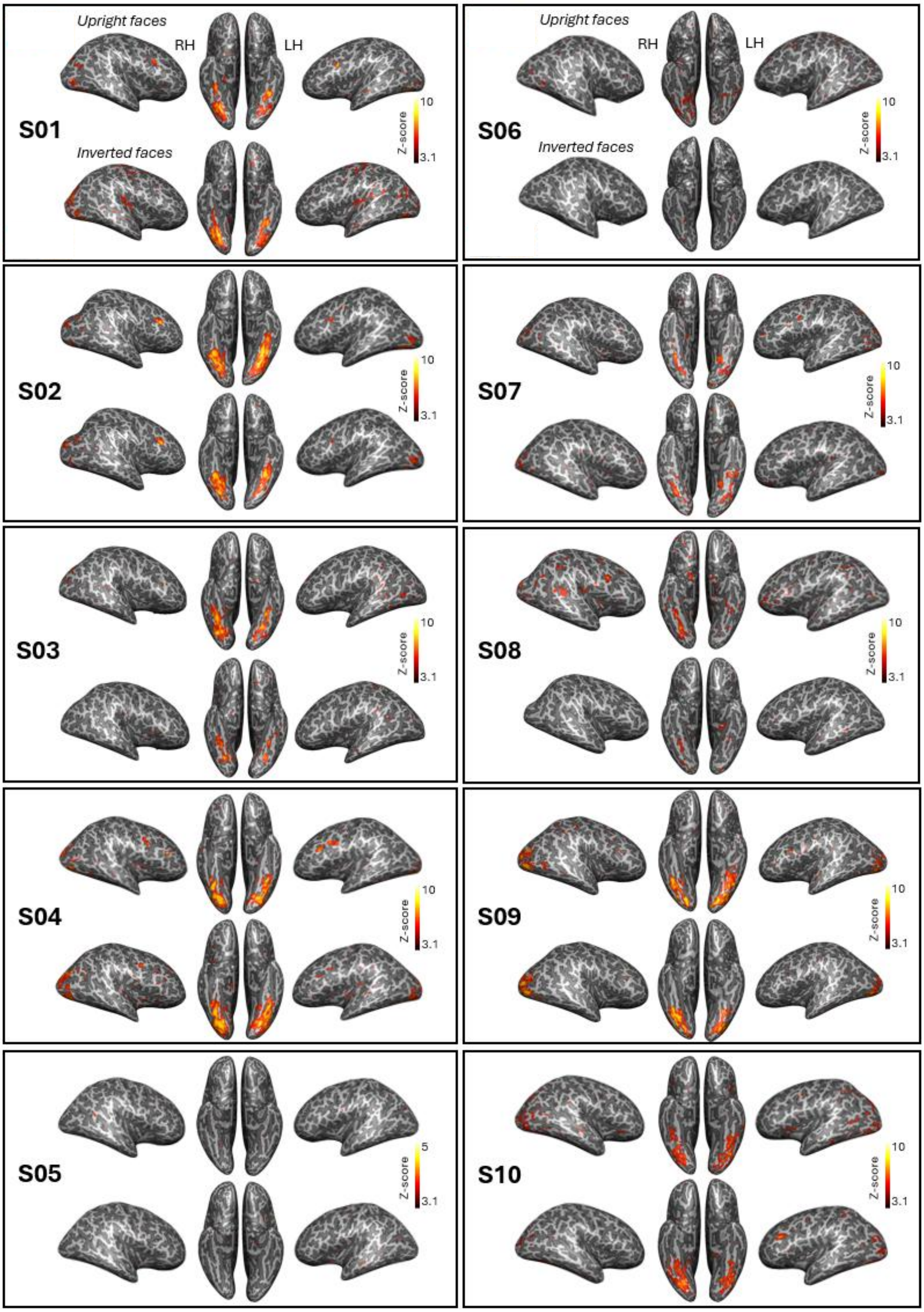


**Supplementary Figure 5. Face individuation responses for upright and inverted faces across high-level changes.** LH: left hemisphere, RH: right hemisphere.

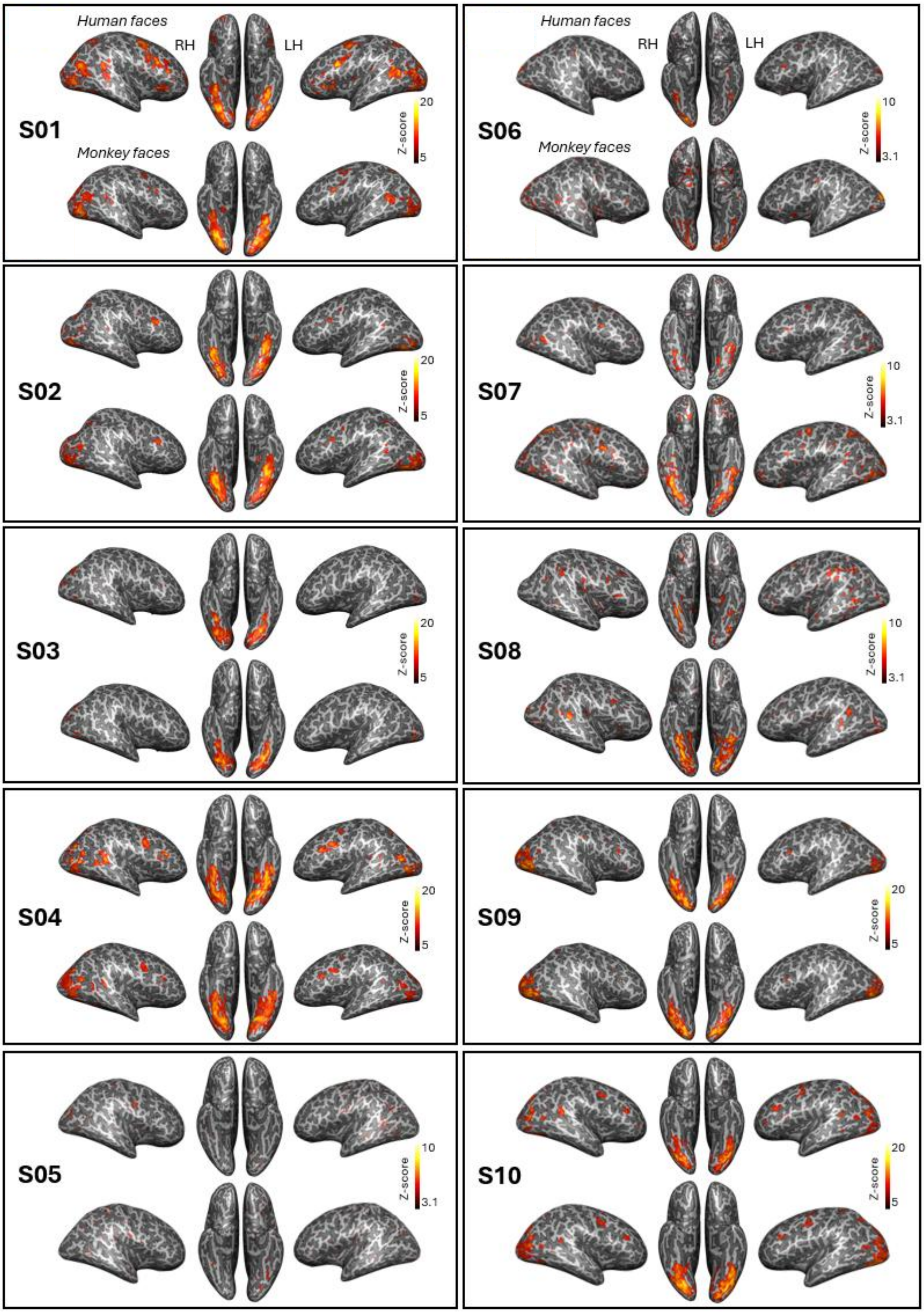


**Supplementary Figure 6. Face individuation responses for unfamiliar human and monkey faces across low-level changes.** LH: left hemisphere, RH: right hemisphere.

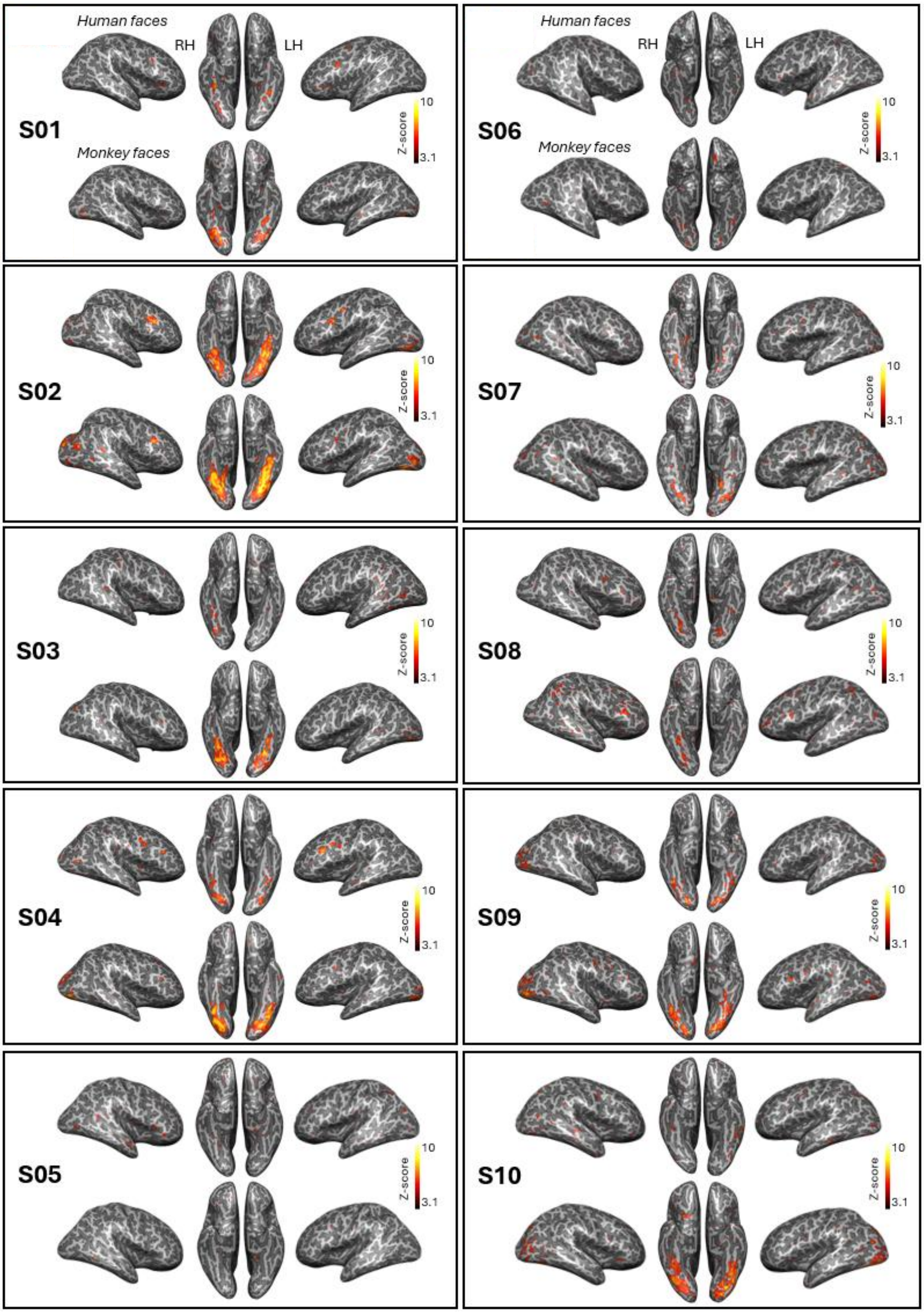


**Supplementary Figure 7. Face individuation responses for unfamiliar human and monkey faces across high-level changes.** LH: left hemisphere, RH: right hemisphere.

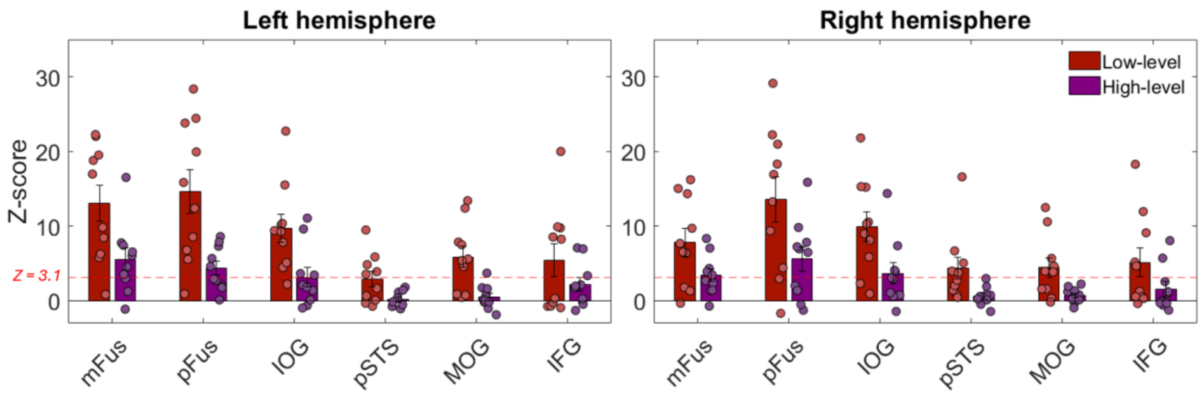


**Supplementary Figure 8. Face individuation responses to unfamiliar faces across low- and high-level image variations in both hemispheres.**

Grouped bars show the mean Z-scores for each face-selective cluster in response to identity changes across low-level (red) and high-level (purple) conditions in the left and right hemispheres. Data are represented as mean ± SEM. The horizontal dotted line represents a threshold level of Z > 3.1, $p < 0.001$, uncorrected; $p < 0.01$ with Bonferroni correction.

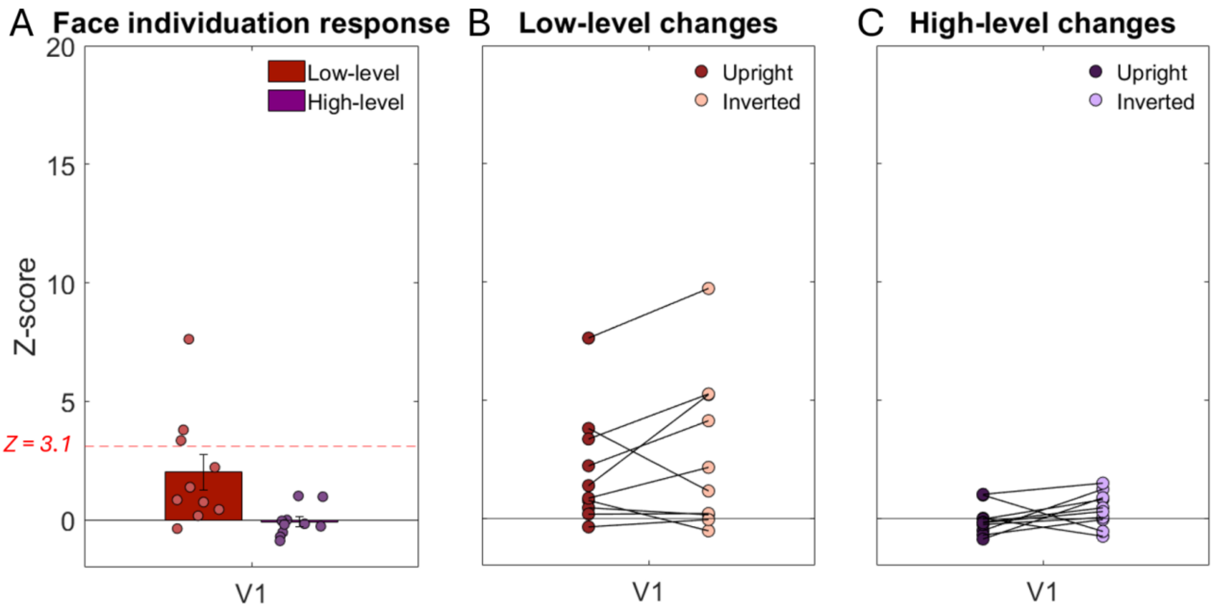


**Supplementary Figure 9. Face individuation responses to upright and inverted faces across low- and high-level image variations in V1.**

**A.** Grouped bars show the mean Z-scores in V1 in response to identity changes across low-level (red) and high-level (purple) conditions. Data are represented as mean ± SEM. The horizontal dotted line represents a threshold level of Z > 3.1, $p < 0.001$, uncorrected; $p < 0.01$ with Bonferroni correction. **B.** For each subject, colored dots provide the mean Z-scores in V1 in response to identity changes, with FIR responses across low-level changes (red: upright faces; light red: inverted faces). **C.** For each subject, FIR responses across high-level changes (purple: upright faces; light purple: inverted faces). The V1 region was defined as the $10^{th}$-order mesh neighborhood around the most posterior point of the calcarine sulcus. *$p < 0.05$; **$p < 0.01$.

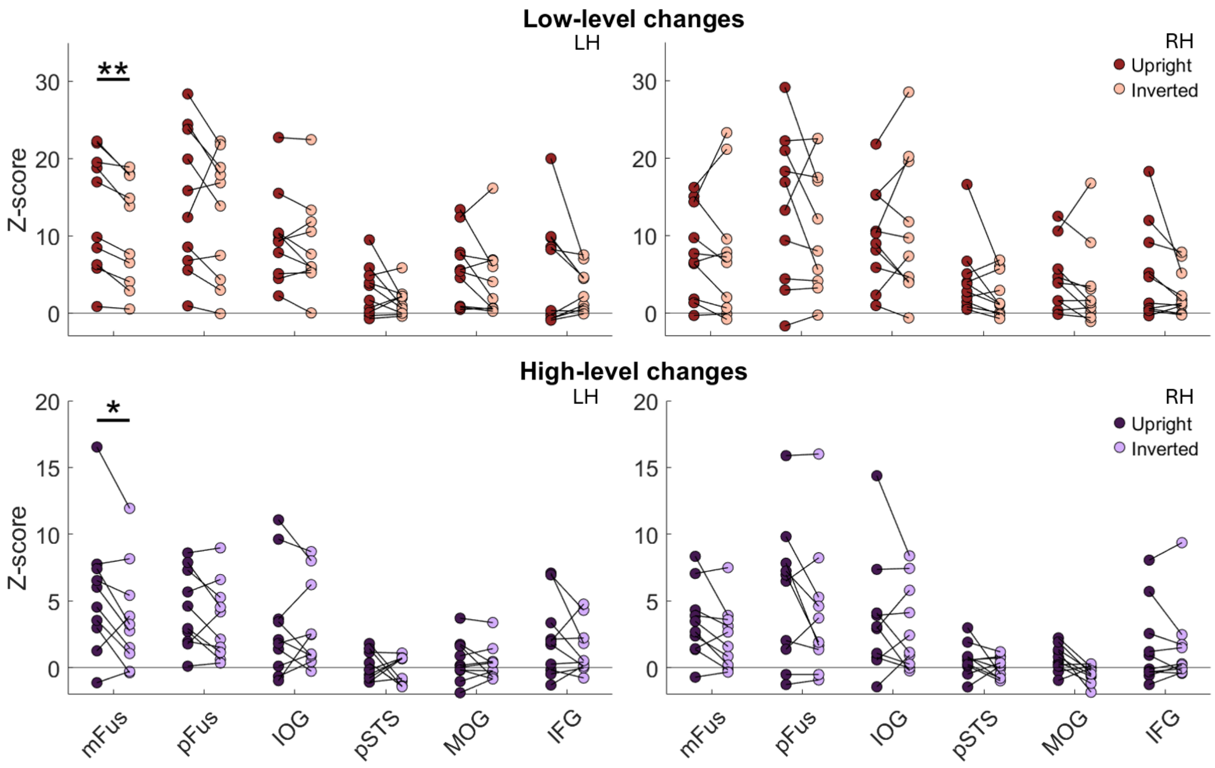


**Supplementary Figure 10. Face individuation responses to upright and inverted faces across low- and high-level image variations in both hemispheres.**
For each subject, colored dots provide the mean Z-scores for each face-selective cluster in response to identity changes, with FIR responses across low-level changes in the upper panel (red: upright faces; light red: inverted faces) and high-level changes in the lower panel (purple: upright faces; light purple: inverted faces). LH: left hemisphere, RH: right hemisphere. **p* < 0.05; ***p* < 0.01.

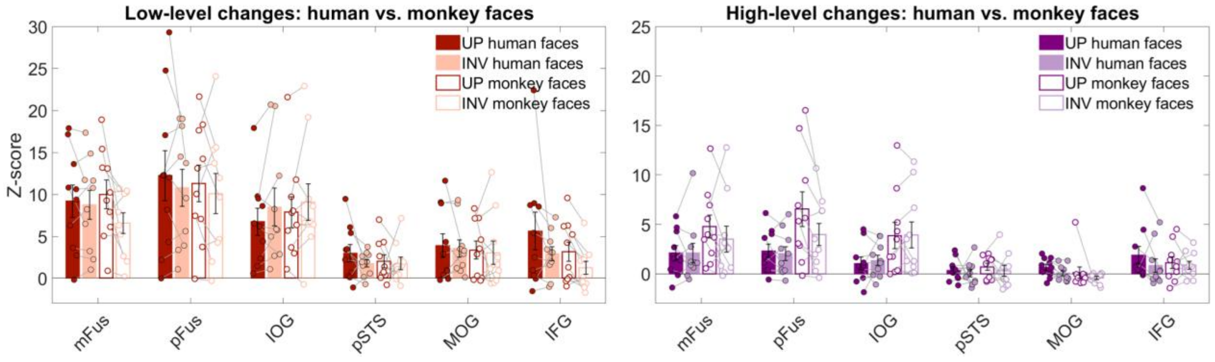


**Supplementary Figure 11. Face individuation responses to upright and inverted faces across low- and high-level image variations for both human and monkey faces.**
Grouped bars show the mean Z-scores for each face-selective cluster in response to identity change, with FIR responses across low-level changes in the left panel (red: upright faces; light red: inverted faces) and high-level changes in the right panel (purple: upright faces; light purple: inverted faces). Solid bars represent human faces, whereas open bars represent monkey faces. Data were averaged across left and right hemispheres. Individual subject data are overlaid.

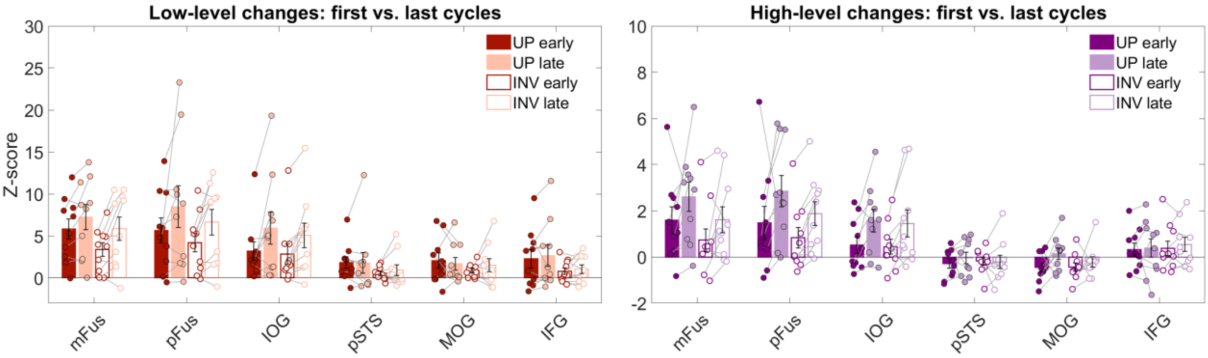


**Supplementary Figure 12. Face individuation responses to upright and inverted faces across low- and high-level image variations for early vs. late cycles.**
Grouped bars show the mean Z-scores for each face-selective cluster in response to identity change, with FIR responses across low-level changes in the left panel and high-level changes in the right panel. Solid bars represent upright faces, whereas open bars represent inverted faces. Data were averaged across left and right hemispheres. Individual subject data are overlaid.

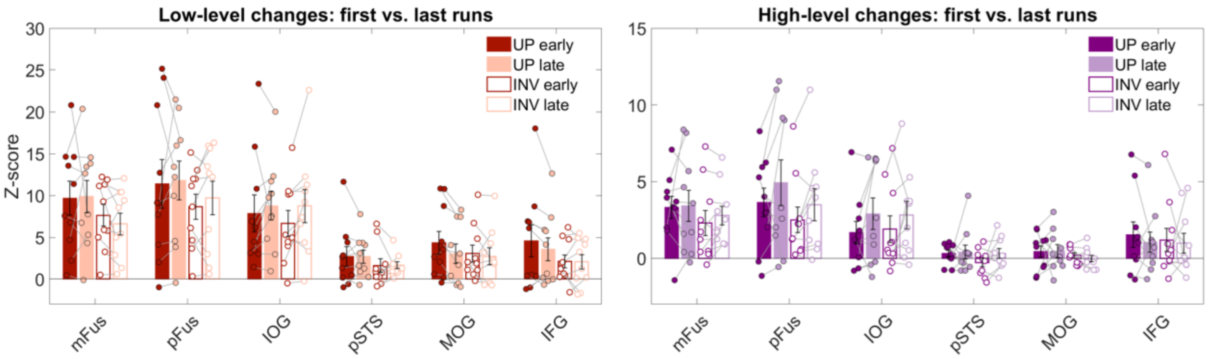


**Supplementary Figure 13. Face individuation responses to upright and inverted faces across low- and high-level image variations for early vs. late runs.**
Grouped bars show the mean Z-scores for each face-selective cluster in response to identity change, with FIR responses across low-level changes in the left panel and high-level changes in the right panel. Solid bars represent upright faces, whereas open bars represent inverted faces. Data were averaged across left and right hemispheres. Individual subject data are overlaid.
